# Optimizing Sequencing Resources in Genotyped Livestock Populations Using Linear Programming

**DOI:** 10.1101/2020.06.29.179093

**Authors:** Hao Cheng, Keyu Xu, Jinghui Li, Kuruvilla Joseph Abraham

## Abstract

**Background:** Low-cost genome-wide single-nucleotide polymorphisms (SNPs) are routinely used in animal breeding programs. Compared to SNP arrays, the use of whole-genome sequence data generated by the next-generation sequencing technologies (NGS) has great potential in livestock populations. However, a large number of animals are required to be sequenced to exploit the full potential of whole-genome sequence data. Thus, novel strategies are desired to allocate sequencing resources in genotyped livestock populations such that the entire population can be sequenced or imputed efficiently.

**Methods:** We present two applications of linear programming models called LPChoose for sequencing resources allocation. The first application is to identify the minimum number of animals for sequencing while meeting the criteria that each haplotype in the population is contained in at least one of the animals selected for sequencing. The second is to sequence a fixed number of animals whose haplotypes include as large a proportion as possible of the haplotypes present in the population given a limited sequencing budget.

**Results:** In both applications LPChoose has similar or better performance than some other methods. The linear programming models we proposed are based on rigorous and well defined optimization techniques and easy and straightforward to implement. LPChoose is available as an open-source package.

## Background

The discovery of genome-wide single-nucleotide polymorphisms (SNPs) and effective ways to assay them has revolutionized genetic analyses of quantitative traits in animal breeding [1, 2, 3, 4, 5]. In conventional breeding programs, low-cost SNP array data are routinely used in genomic selection to estimate breeding values. Compared to SNP arrays, the use of whole-genome sequence data generated by the next-generation sequencing technologies (NGS) has great potential in livestock populations for causal mutation detection [6] in genome-wide association studies and more stable or accurate prediction of breeding values [7] in genomic selection. However, a large number of animals are required to be sequenced to exploit the full potential of whole-genome sequence data. As sequencing is expensive, there is considerable interest in extracting as much information as possible by sequencing a limited number of animals and then imputing to other animals. Several strategies have been proposed to allocate sequencing resources in genotyped livestock populations. Some of these strategies require that certain key individuals be sequenced at high coverage [8] while others consider sequencing large number of animals at lower coverage [9, 10]. The approach to sequence a large number of animals at low coverage offers certain advantages. For example, in [9], the authors comment on the improvement in genotype imputation attained through increasing the number of animals sequenced at low coverage, and in [10], the authors point out the benefits of sequencing many individuals at low coverage in complex trait association studies. The methods we present in this paper are best suited for selecting which animals should be sequenced when selecting a large number of animals at low coverage. The only information used is the haplotypes carried by each animal, and their frequencies. This reliance on the haplotypes present and frequencies to select individuals to sequence is central to two of the methods discussed in [11], AHAP and IWS, which are discussed as well in [12] and in [8]. IWS, for example, makes use of haplotype frequencies to preferentially select animals containing low frequency haplotypes. Another method using haplotype information in [11] is the Genetic Diversity Index, GDI. As the authors of [11] point out, GDI, is quite comparable to IWS. Other methods discussed in these papers rely on pedigree information and will not be discussed here. In addition, no use is made of population histories to decide how many ancestral haplotypes may be present and how this number can affect the choice of animals selected for sequencing [8]. We emphasize that the methods we present assume that a low-density haplotype library has been constructed and this low-density haplotype information on all animals is available. This assumption is also made in [13], [14], and [12]. Furthermore, as in [13, 12], we assume that in order to recover a given haplotype in any animal then at least one animal containing that haplotype must be sequenced. This requirement imposes a number of constraints to be simultaneously satisfied.

The problem of selecting the best set of animals in a population capable of satisfying certain constraints is an optimization problem in a space whose dimension is equal to the total number of animals in the population. In practical applications, and in the examples we will present, the optimization must be performed for tens of thousands of animals subject to tens of thousands of constraints. In general, solving an optimization problem in a space of such high dimension by either numerical or analytical approaches is intractable. However, the optimization problem we are trying to solve here can be addressed through the use of a well-established set of techniques known as linear programming (for an introduction to linear programming and references to original papers see [15]). In linear programming, exact or approximate optimum solutions through numerical approaches can be obtained by using two simplifying features: the objective function to be optimized is linear; the constraints to be satisfied are also linear. Both of these conditions are satisfied for problems we will address in this paper. Linear programming has been previously applied in animal and plant breeding to optimize breeding decisions [16, 17]. However, the use of linear programming for the selection of the most suitable animals to allocate sequencing resources has not been studied. In this paper, we will study the application of linear programming to the allocation of sequencing resources across animals in a population to address two different but related questions which may arise in breeding programs. Both of these questions are closely related to the problem finding the focal individuals described in algorithm 1 of [13].

The first question is to determine the minimum number of animals needed to permit sequence imputation into all other members of the population, and then to identify these animals. Among all animals in the population, initially this subset of animals alone will be subsequently selected for sequencing. The rationale behind initially choosing only these animals for sequencing is as follows: upon sequencing any given animal in this subset, imputation in any other animal carrying the same haplotypes as the sequenced animal can be carried out; this assumption is made in [13]. Thus, once a subset of animals which contains all the haplotypes in the population has been identified, the animals in this subset can be considered to be a starting point for sequencing to eventually permit imputation into the entire population. In order to minimize overall costs, we will attempt to find the smallest subset of animals with this property.

In practice, however, even after the smallest subset of such animals has been identified, it may not be possible to sequence all the animals identified in this manner due to budget constraints. If the budget for sequencing is limited, then it may become necessary to choose a limited subset of animals carrying haplotypes which may be considered to be representative of the population on the basis of being more frequent. Identifying this limited subset of animals is the second question which will be addressed in this paper. The objectives of this paper are to study the use of linear programming models to address both of these questions described above and compare their performance with existing methods.

Much of the motivation for our paper lies in maximizing the haplotype coverage attainable with a fixed number of animals. As we will outline briefly in the discussion section, more efficient coverage could lead to a significant reduction in overall sequencing costs. In this paper, we will present these two applications of linear programming models called LPChoose for sequencing resources allocation. The performance of our LPChoose method will be compared to several approaches including AlphaSeqOpt, IWS, and AHAP. As mentioned before, these applications are related to the issues addressed in algorithm 1 in [13]. We do not discuss issues of haplotype coverage phasing and resolution, which are addressed in algorithm 2 in [13] and in [14]. Some the key ideas in IWS and AHAP for the selection of individuals can be formulated in the language of linear programming. In the results section we will incorporate some concepts central to IWS and AHAP in a linear programming framework and compare the results obtained to those obtained from LPChoose. One important distinction between our implementation of IWS and that in [12] is that we do not restrict our attention to homozygous haplotypes. Both homozygotes and heterozygotes are included in our implementation.

## Methods

Before presenting details of our methods we establish some notations which will be used throughout this paper. For each animal we associate an indicator variable *x*_*j*_ such that *x*_*j*_ ∈ (0, 1) and *j* ∈ (1, 2, …, *n*) where *n* is the total number of animals. *x*_*j*_ = 1(0) means that the animal *j* is selected (not selected) for sequencing. We next introduce binary constants *a*_*ij*_ where *i* ∈ (1, 2, …,, *p*) and *p* is the total number of unique haplotypes. *a*_*ij*_ = 1(0) if animal *j* carries haplotype *i* (or not). Since we are solving systems of inequalities in which the variables are binary valued, we actually work within the framework of integer linear programming, which is more restricted than linear programming. None of the methods we describe make explicit use of a pedigree, but can be applied without modification if a pedigree is known.

The first application of integer linear programming is to identify the minimum number of animals for sequencing while meeting the criteria that each haplotype is contained in at least one of the animals selected for sequencing. If haplotype *i* is carried in at least one animal in the population, then we require that 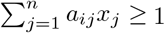 1. This condition can be satisfied only if *x*_*j*_ = 1 for at least one value of 1 ≤ *j* ≤ *n*. A similar constraint must hold separately for all *i* ∈ 1, 2, …,, *p*. Note that all these inequalities can be trivially satisfied by sequencing all animals, i.e., *x*_*j*_ = 1 for all *j* ∈ 1, 2, …,, *n*. This solution almost never leads to the smallest number animals required for sequencing. In order to minimize the number of animals to be sequenced,we additionally require 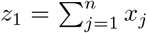, where *z*_1_ is the number of selected animal, to be as small as possible. These inequalities can be collectively written as

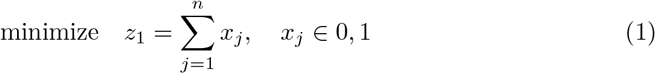

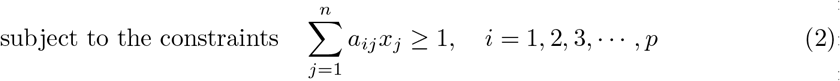

Equation 1 is the objective function to minimize the total number of selected animals. Equation 2 is the set of constraints which ensures that each haplotype is present in at least one of the animals selected for sequencing.

The objective function and the constraints can also be written in matrix notation as

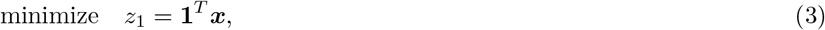

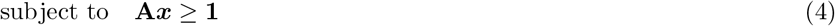

where **A** is the matrix of values *a*_*ij*_ and ***x*** denotes the vector of *x*_*i*_ values. This is a system of inequalities which can be solved by integer linear programming. The solution for *x* will contain some values equal to zero and others equal to one. The *x*_*j*_ which are equal to 1 correspond to the animals which should be sequenced. Once these animals are sequenced, by our earlier assumptions, imputation in the entire population becomes possible.

The second question, to select a fixed number of animals with most common haplotypes, can also be addressed using integer linear programming. In order to prioritize animals with more frequent haplotypes we define a vector **h** with 1 ≤ *i* ≤ *p* whose *i*th element is the frequency of the *i*th haplotype. The vector **h** are then used to define another vector **c** = **h**^*t*^**A**. The number of elements of **c** is equal to the number of animals. Values of **c** are larger for animals which carry many more frequent haplotypes. Thus the values of **c** will be used as a guide to select animals when sequencing resources are limited. In addition, it is important to ensure that the same haplotype is not sequenced in a large number of different animals so as to maximize the haplotype diversity in the animals selected for sequencing. All these different requirements can be summarized in the following set of inequalities

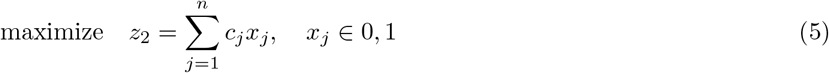

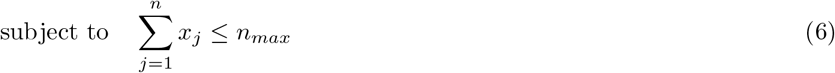

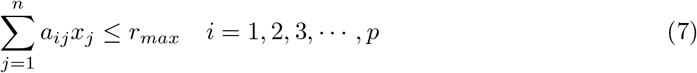

The objective function and the constraints can also be written in matrix notation as

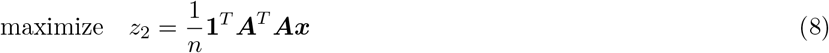

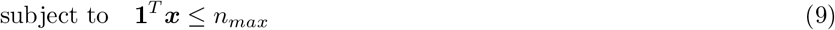

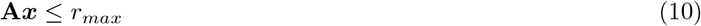

Equation 5 is the objective function to maximize number of most frequent haplotypes represented by the selected animals. Equation 6 is the constraint to set the maximum number of animals to be sequenced *n*_*max*_. Equation 7 is the constraint to ensure that each haplotype covered is at the most covered by *r*_*max*_ animals, where *r*_*max*_ should be a positive integer ≥ 1 and it is chosen to be considerably smaller than *n*_*max*_. Ideally, the objective function is maximized with the value of *r*_*max*_ equal to 1 and the maximization of the objective function in equation 5 is performed in one step. In our data analysis, the value of *r*_*max*_ is set to be 2 to reduce the computational burden. The coefficients *c*_*j*_ in Eq.5 may be freely chosen, as long as the coefficients are positive. With the coefficients chosen as described in [11], it is possible to prioritize the selection of animals in the IWS or the AHAP schemes. A similar choice of coefficients can be made in Eq.3. The highly significant haplotype scheme mentioned in [11] uses coefficients similar to Eq.5, but with some additional multiplicative factors which are not present in our analysis. The IWS scheme we use here is similar to that used in [11], however the greedy approximation used in [11] is different from the approximation we use.

Linear programming problems described above cannot be solved analytically. However, they can be solved using branch and bound methods [18], which are guaranteed to converge to the global optimum. In practice, for reasons which will be discussed later, the convergence can be very slow. Thus we will use a fast approximation in the second application to select a fixed number of animals. Instead of selecting *n*_*max*_ animals at once, we will first select 2 animals (i.e., *n*_*max*_ = 2). Once the first two animals have been identified, these animals and all the haplotypes present in them are removed. Then the optimization in equations 5, 6, and 7 is repeated but with the objective function in equation 5 suitably modified to reflect the absence of the first two animals. This procedure can be carried out until the desired number of animals has been found. A numerical example is shown in the Appendix to demonstrate the use of linear programming in both applications.

### Data analysis

Genotype data for five different scenarios were simulated using AlphaSimR [19]. The number of generations considered in these 5 scenarios are 5, 10, 15, 30, or 50. These scenarios resembled modern cattle populations. The genome of 10,000 segregating loci on 10 chromosomes was simulated using the “cattle genome” option in AlphaSimR. A quantitative trait controlled by 150 QTL of effects sampled from standard normal distributions distributed equally on 10 chromosomes was simulated. First, founders of 1,000 cattle of equal sex ratio were generated. At each generation, the best 25 males were selected as sires on the basis of their highest breeding value and mated to all 500 females as dams to produce next generations with 1000 cattle of equal sex ratio. In our analysis, 10 replicates were simulated for each of the five scenarios. From the above, all individuals had haplotypes for 10,000 SNPs distributed equally across the 10 chromosomes. In each population, haplotype blocks of length 100 SNPs were obtained across 10 chromosomes. A mismatch of up to 10% was used to ensure that haplotypes with small differences were considered as identical. In the results which will be presented, the selection of a fixed number animals will also be made after the exclusion of haplotypes with a frequency of less than 1%, similar to the strategy adopted in [20]. The results in [12] are also based on the exclusion of low frequency haplotypes but with a more drastic restriction on haplotype frequency than in [20]. A brief description of the simulation scenarios is presented in table 1. An open-source, publicly available Julia package called LPChoose (https://github.com/reworkhow/LPChoose.jl) has been developed. A publicly available implementation of linear programming, the GNU Linear Programming Kit (GLPK), is used in LPChoose. In particular, for the sake of the reproducibility of our results, we do not make use either directly or indirectly of any proprietary solvers for linear programming problems.

**Table 1.** Simulation scenarios.

| scenario | total number of animals | number of unique haplotypes | number of unique common haplotypes |
| --- | --- | --- | --- |
| 1 | 6,000 | 27,399 ± 332 | 2,638 ± 42 |
| 2 | 11,000 | 37,922 ± 758 | 2,376 ± 50 |
| 3 | 16,000 | 44,591 ± 826 | 2,216 ± 49 |
| 4 | 31,000 | 56,036 ± 1,313 | 1,959 ± 54 |
| 5 | 51,000 | 61,418 ± 2,023 | 1,815 ± 44 |

## Results

The minimum numbers of animals identified in LPChoose, AlphaSeqOpt, IWS, and AHAP to contain all unique haploptypes in five populations are shown in table 2. The minimum numbers of animals identified by LPChoose were consistently smaller than those from AlphaSeqOpt, IWS, and AHAP. As AlphaSeqOpt does not directly output the minimum number of animals needed to cover all haplotypes, the values in the last column of table 2 were obtained from repeated runs of AlphaSeqOpt with varying numbers of animals to be selected. The results of table 2 suggest that the difference among these methods in the smallest number of animals required grows as the size of the population increases, especially between LPChoose and AlphaSeqOpt. We emphasize that the results in column 1 of table 2 obtained from linear programming are obtained without any approximations, and are thus equal to the theoretically lowest values attainable for this kind of problem. It is important to note that, when LPChoose is used to select the minimum number of animals to cover all haplotypes, some haplotypes may be covered by multiple animals. This introduces a certain level of redundancy in sequencing which may be desirable for example when certain haplotypes should be covered more frequently. This point is discussed in detail in [14], where the authors assign a score function for haplotypes which reflects the target haplotype coverage and the number of times a given haplotype is included among all the animals assigned for sequencing.

**Table 2.** Minimum number of animals identified representing all haplotypes in the population.

| scenario | minimum number of animals (mean ± SD) |  |  |  |
| --- | --- | --- | --- | --- |
|  | LPChoose | AlphaSeqOpt | IWS | AHAP |
| 1 | 4,086 ± 13 | 4,988 ± 30 | 4,154 ± 30 | 5,998 ± 2 |
| 2 | 6,663 ± 18 | 8,563 ± 51 | 6,802 ± 84 | 10,999 ± 2 |
| 3 | 8,648 ± 34 | 11,563 ± 132 | 8,821 ± 75 | 15,998 ± 3 |
| 4 | 12,229 ± 86 | 17,506 ± 275 | 12,738 ± 225 | 30,997 ± 6 |
| 5 | 14,241 ± 119 | 21,098 ± 494 | 15,028 ± 356 | 50,999 ± 1 |

LPChoose, AlphaSeqOpt, IWS, and AHAP were also used to select a fixed number of animals (100 animals) whose haplotypes represent maximum proportions of the total haplotypes in the population. The results obtained are shown in tables 3 and 4. In table 3, the proportion of all haplotypes represented by 100 selected animals were compared. Animals identified in LPChoose consistently covered higher proportion of all haplotypes than those in AlphaSeqOpt and AHAP but lower than IWS. Only a small proportion (0.1-0.3) of all haplotypes were covered in all three methods. In table 4, the proportion of common haplotypes represented by 100 selected animals were compared. LPChoose performed better than other three methods consistently across different populations, and the proportions of common haplotypes identified in LPChoose were usually higher than 0.99, followed by IWS with slightly lower proportions and those in AlphaSeqOpt are about 0.8-0.9.

**Table 3.** Proportion of all haplotypes represented by 100 selected animals in the population.

| scenario | proportion of all haplotypes (mean ± SD) | | | |
| --- | --- | --- | --- | --- |
|  | LPChoose | AlphaSeqOpt | IWS | AHAP |
| 1 | 0.2218 ± 0.0036 | 0.2052 ± 0.0048 | 0.2504 ± 0.0032 | 0.1441 ± 0.0041 |
| 2 | 0.1700 ± 0.0020 | 0.1528 ± 0.0048 | 0.1951 ± 0.0024 | 0.1007 ± 0.0024 |
| 3 | 0.1498 ± 0.0026 | 0.1319 ± 0.0047 | 0.1735 ± 0.0009 | 0.0805 ± 0.0029 |
| 4 | 0.1249 ± 0.0032 | 0.1119 ± 0.0037 | 0.1464 ± 0.0022 | 0.0584 ± 0.0024 |
| 5 | 0.1145 ± 0.0027 | 0.1084 ± 0.0036 | 0.1370 ± 0.0023 | 0.0435 ± 0.0025 |

**Table 4.** Proportion of common haplotypes represented by 100 selected animals in the population.

| scenario | proportion of common haplotypes (mean ± SD) | | | |
| --- | --- | --- | --- | --- |
|  | LPChoose | AlphaSeqOpt | IWS | AHAP |
| 1 | 0.9934 ± 0.0017 | 0.9239 ± 0.0032 | 0.9917 ± 0.0017 | 0.8520 ± 0.0159 |
| 2 | 0.9911 ± 0.0028 | 0.9041 ± 0.0071 | 0.9899 ± 0.0017 | 0.8288 ± 0.0142 |
| 3 | 0.9921 ± 0.0018 | 0.8766 ± 0.0171 | 0.9875 ± 0.0032 | 0.7625 ± 0.0257 |
| 4 | 0.9899 ± 0.0021 | 0.8444 ± 0.0109 | 0.9780 ± 0.0030 | 0.7004 ± 0.0247 |
| 5 | 0.9993 ± 0.0006 | 0.8335 ± 0.0088 | 0.9600 ± 0.0061 | 0.6287 ± 0.0336 |

Table 3 shows that IWS performs better than LPChoose in terms of unique haplotype coverage in the absence of a lower limit on haplotype frequencies. IWS preferentially selects animals with low-frequency haplotypes by assigning larger weights to individuals containing rare haplotypes [12], which is a hallmark of IWS unlike LPChoose where more common haplotypes are prefered. IWS, however, can be easily accommodated in LPChoose by changing the weights in equation 3 and setting *n*_*max*_ = 1 and *r*_*max*_ = 1. Thus it can be accommodated in a linear programming framework and considered as a version of LPChoose but with modified weights. The authors of IWS mention that even after selecting animals based on the analysis of haplotypes which appear above a certain frequency threshold, the selected animals also contain haplotypes with a frequency below that threshold. This is also the case when animals are selected in LPChoose using equations 8,9, and 10.

## Discussion

In this paper, linear programming models are used to answer two questions for the allocation of sequencing resources. The first question addressed is to identify the minimum number of animals whose haplotypes represent all the haplotypes in the population. The second question we address is to choose for sequencing a fixed number of animals whose haplotypes represent the maximum proportion of the total or common haplotypes in the population. In practice, the second question usually comes into play due to the limited sequencing budget. Because of restrictions in the sequencing budget, it may be possible to sequence only a fixed number of animals which may turn out to be less than the number of animals needed to cover all the haplotypes in the population. In this scenario, the animals selected should be chosen based on some additional criteria, for example, the animals selected should carry as large a proportion as possible of the haplotypes in the population.

In the results presented, we have assumed that in order to impute any given haplotype in any animal that haplotype must be present in at least one animal selected for sequencing. In addition, it has been implicitly assumed that once an animal has been sequenced then imputation on all other animals carrying the haplotypes of the sequenced animals can potentially be carried out. Indeed, the structure of the linear programming equations and the resulting solutions rely on this first assumption. The results we present suggest our methods offer an improvement over those previously published using similar criteria by way of requiring the sequencing of fewer animals. This economy in the terms of the number of animals needed may offer very significant advantages in terms of costs of imputation. In [21], the authors discuss the use of hybrid peeling for imputation and suggest that for the phasing of haplotypes in each animal initially selected for sequencing, six additional animals, the parents and the grandparents should also be sequenced at some coverage. If the number of animals initially selected for sequencing can be minimized without sacrificing haplotype coverage, then the number of additional animals which have to be sequenced could potentially be significantly reduced leading to a sizeable economy in sequencing costs. As in the case of [22], the approach in [21] makes extensive use of pedigree data. Even though LPChoose makes no explicit use of the pedigree, LPChoose as described in the methods section can be used in conjunction with pedigree based methods such as hybrid peeling provided the haplotypes are known. Other criteria have been used to determine the best animals for sequencing, including the expected accuracy of genotype imputation [21, 23]. It has been suggested, based on extensive simulations that genotype imputation accuracy can be improved when a larger number of animals are sequenced at lower coverage as opposed to a smaller number of animals at high coverage (assuming the product of read depth and number of sequenced animals is fixed) [9]. Further advantages of sequencing large number of animals at low coverage have also been discussed by [8]. As our results indicate that for a fixed number of animals, LPChoose covers more haplotypes than competing methods, the methods in this paper could also be used as a guide to select animals which permit a higher level of genotype imputation accuracy.

It is interesting to note that in [12], the authors suggest that *“a sequencing project that targets all observed haplotypes is likely to be prohibitively expensive and never-ending”*. Our results for the first application suggest that this may not be the case if linear programming methods are used. Although including all haplotypes, even those which appear in a single animal increases the size of the final set of animals selected, the inclusion of these very rare haplotypes facilitates the rapid identification of which animals must be included in the final result. Furthermore, if these animals also carry certain higher frequency haplotypes, then these haplotypes can be accounted for very early in the analysis further simplifying the problem. In the same paper the authors point out that the accuracy of imputation of rare variants is affected by the lower limit put on haplotype frequencies and suggest the importance of including lower frequency haplotypes; the methods presented in this paper are well suited for the inclusion of low frequency haplotypes.

There are some unavoidable limitations associated with the linear programming methods we have used. The presence of these limitations has already been suggested earlier, e.g., the results presented in table 4 are based on an approximate solution to the linear programming problem. To get additional insight into the technical difficulties which necessitate the use of approximations we will rephrase the problem of identifying the smallest set of animals which carry all the haplotypes in the population. We begin by defining a network consisting of the animals and haplotypes as follows. For each animal we define a vertex in an undirected graph and for each haplotype we also define a vertex in the same undirected graph. The edges in this undirected graph are defined as follows: there are no edges between vertices which define haplotypes; there are also no edges between vertices which represent animals.

There is an edge between two vertices if one of the vertices represents an animal, the other represents a haplotype, and the animal contains the haplotype. This graph is thus a bipartite graph, with two subsets of vertices, representing animals and haplotypes, and the only edges present are those between the two subsets.

The methods presented earlier in this paper for finding the minimal number of animals containing all the haplotypes in the population are essentially the solution to a set cover problem on a bipartite graph using integer linear programming, which is a very well established method for the solution of the set cover problem in a general context [24]. However, there is no known polynomial time algorithm for solving the set cover problem. Thus as the number of haplotypes grows, computational costs associated with an exact solution may become prohibitive. The graphical formulation we use is that of an undirected graphical model, a related but different formalism, directed graphical models, have been used for IBD detection on the Beagle Suite of Programs [25]

The second computational issue we discuss, that of finding the best small sub-set of animals, can also be interpreted in terms of a set cover problem with additional weighting factors corresponding to haplotype frequencies. In light of the correspondence of the questions we address with a class of problems in combinatorial optimization for which there is no known efficient algorithm, it is not surprising that approximations were required. Much of the difficulty arises due to the discrete nature of the *x* variables. In previous papers using continuously varying genomic fractions to optimize mating decisions to control for inbreeding [26, 27] this difficulty does not arise.

The approximations we have made use of are not the only approximations which can be employed in problems of this nature. Another class of approximation which can be easily incorporated into the framework of linear programming is to relax the requirement in equation 3 and equation 4 that the *x* variables be binary. Instead they can be set to be continuous in the range from 0 to 1. This relaxation transforms the original problem to a problem can be solved in polynomial time. However, after relaxation the solutions for the *x* variables may lie between 0 and 1. One possibility to decide which animals should be discarded and which should be sequenced is to choose a threshold (e.g., 0.8) such that all *x* values above the threshold are set equal to one and all values below the threshold are set equal to zero. In this way animals to be sequenced are identified. However, there is no guarantee that with this thresholding scheme the animals to be sequenced will contain all the haplotypes. If there are some haplotypes that have not been covered, then the additional animals to be sequenced can be identified by solving the relaxed versions of equation 3 and 4 subject to the additional requirements that the *x* variables for the animals found in the previous approximation be conditioned to 1. With this conditioning requirement, many of the constraints in equation 4 are now satisfied. With many constraints satisfied, Equation 3 and 4 can be solved again more easily. This cycle can be repeated until all constraints are satisfied. Once a sufficient number of constraints are satisfied, it may be that there is no longer any need to address the relaxed form of the equations and the remaining constraints be solved using binary *x* values. The scheme outlined offers a way to get approximate solutions for equation 3 and 4 when the size of the data set makes an exact solution computationally unfeasible.

Despite these fundamental limitations necessitating the use of approximations, linear programming methods offer many advantages. For example, with linear programming multiple solutions can be identified if needed. Another example of the flexibility of linear programming methods arises if one animal must be included in the set of animals to be sequenced. In this case that animal can be included by an additional constraint in equation 4, and the other animals selected in the most economical way after taking into account haplotypes present in the animal included by requirement. The requirement that multiple pre-selected animals have to be included can also be accommodated by the inclusion of multiple additional constraints.

As we have mentioned earlier even when minimizing the number of animals sequenced there is some redundancy in haplotype coverage. If this level of redundancy is not adequate for haplotype resolution, then additional redundancy can be imposed by setting certain values on the right hand side of Eq.4 to be larger than 1. In this case additional animals may be required to be sequenced, however the total number of animals is still minimized provided no approximations are made in obtaining the modified versions of the solution of Equations 2.

Finally, the methods we have developed are motivated by the situation in which sequencing coverage is low with no animals selected for larger fold coverage. If however, certain animals are sequenced to higher coverage, the effect on haplotype coverage can be roughly approximated by multiplying the appropriate columns of the **A** matrix by the higher fold coverage and repeating the previous analyses.

## Conclusions

We have illustrated the use of linear programming for optimizing the allocation of sequencing resources. The results presented show that linear programming offers certain advantages over some currently known approaches. The methodology we have illustrated here could be used in conjunction with methods for phasing and haplotype resolution to maximally optimize the use of limited sequencing budgets.

## Appendix

### Numerical example

A numerical example is shown below with haplotype information for 5 animals and 2 haplotypes blocks (table 5).

**Table 5.** haplotypes at 2 blocks for 5 individuals used in the numerical example.

| Animals | block 1 | block 2 |
| --- | --- | --- |
| 1 | 1 2 | 7 8 |
| 2 | 2 3 | 9 9 |
| 3 | 3 4 | 10 11 |
| 4 | 2 4 | 11 12 |
| 5 | 5 6 | 12 13 |

In table 5 numerical labels for haplotpyes should be interpreted as purely qualitative labels. As there are 5 animals and 13 distinct haplotypes, the corresponding **A** matrix has 5 columns and 13 haplotypes and is shown below.

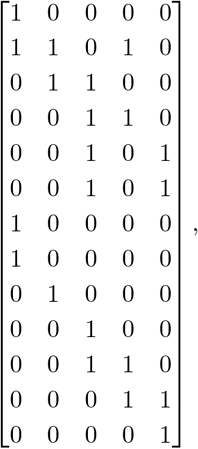

where the element from its *i*th row and *j*th column is *a*_*ij*_, and *a*_*ij*_ is a binary variable such that *a*_*ij*_ = 1(0) if animal *j* contains haplotype *i* (or not).

The first application of integer linear programming to select the minimum number of animals to identify all haplotypes is formulated as

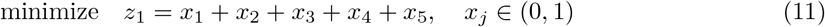

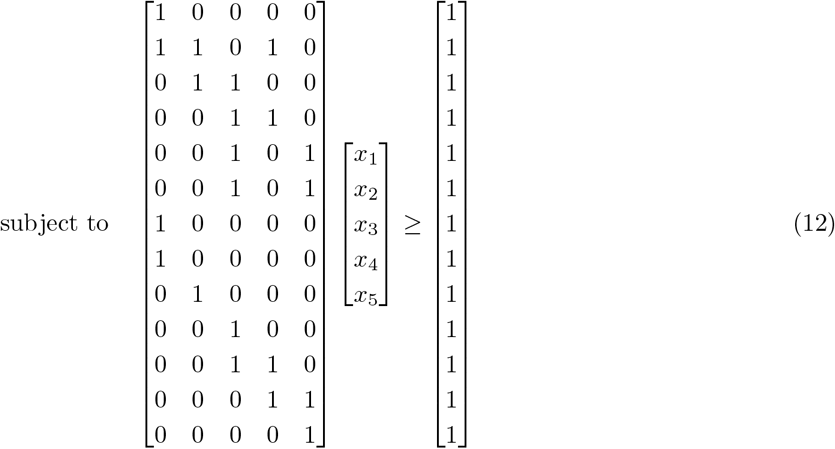

As the result, animal 1,2, 3, and 5 are selected to be sequenced.

The second application of integer linear programming to select a fixed number of animals, e.g., 2 animals, with most common haplotypes is formulated as

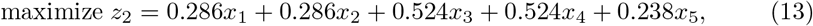

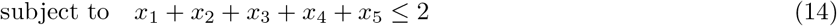

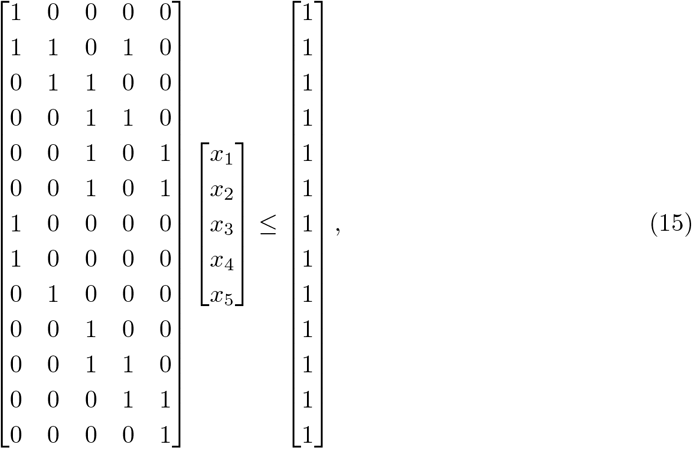

At the end, animal 1 and animal 3 are selected.

## Author’s contributions

KJA, HC conceived the study. KJA, HC, KX, JL contributed to the development of the methodology. KJA, HC, KX, JL developed the LPChoose package. KX, JL preformed the simulation and analysis. HC, KJA wrote the manuscript. All authors read and approved the final manuscript.

## Acknowledgements

This study was financed in part by the Coordenação de Aperfeiçoamento de Pessoal de Nível Superior Brasil(CAPES)-Finance Code 001.

